# Kv1 potassium channels control action potential firing of putative GABAergic deep cerebellar nuclear neurons

**DOI:** 10.1101/779082

**Authors:** Jessica Abigail Feria Pliego, Christine M. Pedroarena

**Affiliations:** Graduate School of Cellular and Molecular Neurosciences, University of Tübingen; Department for Cognitive Neurology, Hertie-Institute for Clinical Brain Research, 72076 Tübingen, Germany; Systems Neurophysiology, Werner Reichardt Center for Integrative Neuroscience, University Tübingen, 72076 Tübingen, Germany

**Keywords:** cerebellum, intrinsic properties, EAT-1, KCNA1, KCNA2, Dendrotoxin

## Abstract

The Kv1 voltage-gated potassium channels (kv1.1-1.8) display characteristic low-threshold activation ranges what enables their role in regulating diverse aspects of neuronal function, such as the action potential (AP) threshold and waveform, and thereby influence neuronal excitability or synaptic transmission. Kv1 channels are highly expressed in the cerebellar cortex and nuclei and mutations of human Kv1 genes are associated to episodic forms of ataxia (EAT-1). Besides the well-established role of Kv1 channels in regulating the basket-Purkinje cells inhibitory synapses of cerebellar cortex, cerebellar Kv1 channels regulate the principal deep cerebellar nuclear neurons activity (DCNs). DCNs however, include as well different groups of GABAergic cells that project locally to target principal DCNs, or to the inferior-olive or recurrently to the cerebellar cortex, but whether their function is controlled by Kv1 channels remains unclear. Here, using cerebellar slices from the GAD67-GFP line mice to identify putative GABAergic-DCNs and specific Kv1 channel blockers (dendrotoxins-alpha/I/K (DTXs)) we provide evidence that putative GABAergic-DCNs spontaneous and evoked activity is controlled by Kv1 currents. DTXs shifted in the hyperpolarizing direction the voltage threshold of spontaneous APs in GABAergic-DCNs, increased GABAergic-DCNs spontaneous firing rate and decreased these neurons ability to fire repetitively action potentials at high frequency. Moreover, in spontaneously silent putative nucleo-cortical DCNs, DTXs application induced depolarization and tonic firing. These results strongly suggest that Kv1 channels regulate GABAergic-DCNs activity and thereby can control previously unrecognized aspects of cerebellar function.

## INTRODUCTION

The Shaker-related Kv1 (Kv1.1-Kv1.8)channels display characteristic low threshold voltage gated potassium currents, well suited to regulate action potential (AP) threshold and waveform and consequently global or local excitability, axonal and synaptic transmission ^1, 2, 3, 4^, as well as the timing, pattern, and precision with which neurons generate spikes ^5, 6^. Kv1 channel proteins (except for the Kv1.7 member) are expressed in the brain, where native channels are usually composed by heteromeric, but also homomeric assembly of four Kv1-alpha subunits potentially associated to different kv1-beta subunits. The composition, clustering, and targeting to different cellular compartments is assumed to support diverse functional roles ^2 4^. The discovery of toxins specific for Kv1 channel subtypes has been crucial to elucidate Kv1 functional roles in different brain areas and neurons ^7^.

The finding of point mutations of the gene KCNA1 (Kv1.1) associated to a familiar disorder characterized by attacks of ataxia with or without myokimia, epidosodic ataxia type-1 (EAT-1), ^8^ indicated a critical role for Kv1.1 channels in cerebellar function. Later, it was found that mutations in the KCNA2 gene (Kv1.2) may result in mice ^9^ and humans in ataxia and convulsions ^10, 11, 12^. Kv1 channels, in particular, Kv1.1, 1.2 and Kv1.6 are highly expressed in the cerebellar cortex, and in particular Kv1.1 and Kv1.2 channels are highly expressed in the terminals of the basket cell interneurons of the cerebellar cortex ^13, 14^. Consistently, application of a specific Kv1 channel blocker, dendrotoxin-alpha (DTX_alpha)^7^, greatly enhances the amplitude and frequency of spontaneous inhibitory postsynaptic potentials (IPSCs) in Purkinje cells, likely mediated by the block of a fraction of the synaptic terminal potassium outward currents ^15^. Furthermore the frequency and amplitude of spontaneous IPSCs in Purkinje cells is found enhanced in a mouse model of EAT-1^16^. Therefore, aberrant evoked synaptic transmission at the basket-Purkinje cell synapses is a likely major cause of the ataxic symptoms of EAT-1 ^4^. However, since Kv1, particularly kv1.1 and Kv1.2 channels are also expressed in the deep cerebellar nuclei neurons (DCN), apparently forming heteromers, it has been suggested that DCNS aberrant Kv1 channels could contribute to the EAT-1 cerebellar symptomatology ^17^, supported by findings that the activity of putative glutamatergic and thalamic - projecting DCNs is sensitive to Kv1 channel blockers ^17^. In addition to the principal DCNs, the cerebellar nuclei are composed by diverse groups of GABAergic neurons which are assumed to play important roles in cerebellar function but it remains unclear whether and how their function is regulated by Kv1 channels as is the case for the principal DCNs. The GABAergic DCNs include: first, the group of small GABAergic neurons forming the nucleo-olivary pathway, which is thought that by directly inhibiting and/or regulating the electrical coupling of inferior-olive cells control the spatio-temporal pattern and probability of cerebellar complex-spikes occurrences ^18, 19, 20, 21, 22, 23^. Second, a group of local interneurons co-releasing GABA and glycine that innervates the principal DCNs and could, therefore, modulate the cerebellar output ^24, 25, 26^. And, third, a group of medium sized GABAergic/glycinergic cells, mostly silent in slices in contrast to all other DCNs, originating a feedback inhibitory pathway to the cerebellar cortex that targets Golgi cells, potentially regulating activity at the cerebellar cortex input stage ^26, 27^. Here, using cerebellar slices from GFP -GAD67 mice line to identify putative GABAergic-DCNs and a pharmacological approach we found evidence that indeed, the activity of GABAergic-DCNs is regulated by Kv1 channels and thus their alteration could contribute to the symptomatology of EAT-1.

## RESULTS

Here, to elucidate whether Kv1 channels control the activity of GABAergic deep cerebellar nuclear neurons (DCNs), we used cerebellar slices from the mature (>P21) GAD-GFP mice ^28^, their wild type littermates, or mice with the same background to obtain whole cell current clamp recordings of DCNs from the lateral or interpositus nuclei before and during application of pharmacological blockers of Kv1 channels (see later for details). To achieve conditions close to those found *in situ* the slices were maintained during the recordings at close to physiological temperature (33-35°C) and were perfused with ACSF containing physiological extracellular calcium concentrations (1.5 mM).

GFP fluorescence was used to identify putatively GABAergic DCNs ^28^ (GAD+DCNs). Non-GFP fluorescent medium sized DCNs (diameter >18μM) recorded from the GAD-GFP mice line slices, and large DCNs (diameter > 21μm) from any mouse line displaying spontaneous spiking activity were identified as putatively glutamatergic principal DCNs ^28^.

In a first series of experiments we analyzed whether dendrotoxin-alpha (DTX-alpha, 100nM), which blocks with low EC_50_ channels containing at least one Kv1.1, Kv1.2 and Kv1.6 subunit ^2, 7^, modulates the spontaneous action potentials (APs) and firing properties of GAD+DCNs. The toxins were applied under the presence of neurotransmitter blockers (see methods), to prevent indirect effects due to changes in the spontaneous release of excitatory or inhibitory neurotransmitters and the effect was analyzed after approximately 10 minutes to assure a steady state level was reached (Figs. 1 and 2). The most consistent effect induced by the DTX-alpha application was a shift in the voltage threshold of spontaneous APs in the hyperpolarizing direction. Because DTX application often induced a change in membrane potential during the inter-spike interval (ISI) (see later, and inset Fig 1A), to be able to evaluate whether DTX directly affected the spike threshold after a period of recording at the spontaneous membrane potential, we injected constant current to bring the ISI membrane potential close to control levels (Fig. 1A). This correction is necessary because in spontaneous spiking DCNs sodium channels are partially inactivated between spikes, rendering the spike waveform extremely sensitive to changes in the ISI membrane potential (ISI-Vm)^29, 30^. The analysis of action potential waveform was then carried in averaged action potentials with inter-spike membrane potential matching that in control conditions. To detect the threshold we used phase plots as previously (Fig. 1B, see methods for details) ^30^. The threshold hyperpolarized in all 6 neurons investigated and in average the change in threshold was −1.4 mV ± 0.26 (paired t-test, P= 0.003, n=6, Fig. 1C, top left panel). Interestingly, the hyperpolarizing shift in spike voltage threshold occurred despite associated increases in spontaneous firing rate and decreased spike amplitude, suggesting a strong control by Kv1 channels of this parameter, as will be discussed later.

**Figure 1.**
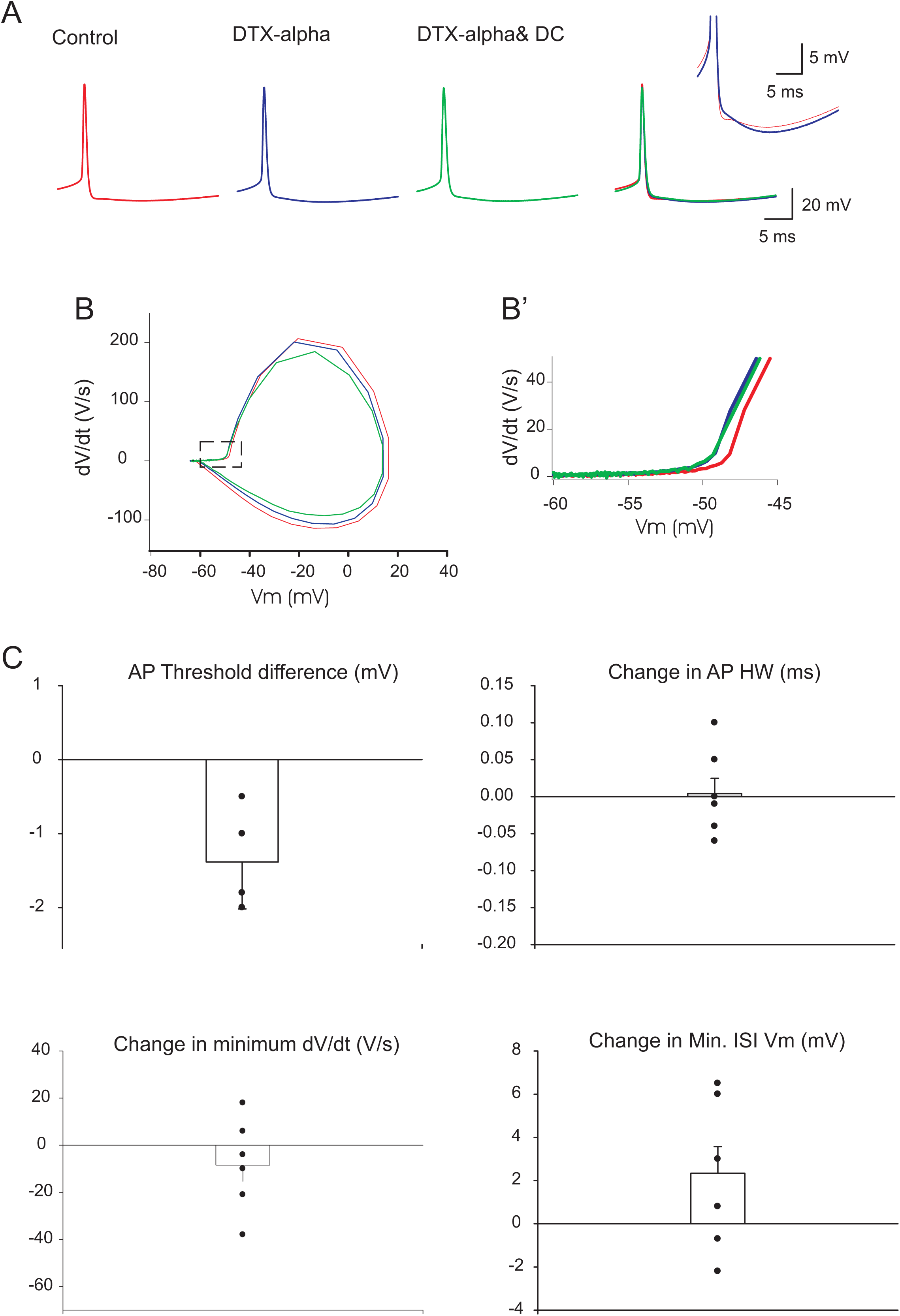
The action potential waveform of GAD+DCNs is sensitive to DTX-alpha application. A- From left to right: Typical example of averaged spontaneous action potentials recorded from one GAD+DCN during application of NT blockers (control), during DTX-alpha (100nM) application (DTX-alpha), and after correcting the ISI Vm by current injection during DTX-alpha application (DTX-alpha& DC), and all traces superimposed. The inset depicts in expanded scale the control and DTX-alpha traces. B- Phase plots corresponding to the traces in A (the derivative of the Vm as a function of the Vm). B’-Detail of B in expanded time scale to show the hyperpolarizing shift in the rapid raise in dV/dt signaling the AP threshold. C- Mean change in AP threshold for all recorded GAD+DCNs under DTX-alpha and individual results superimposed (some data points overlap). D- Mean change in AP HW for all recorded GAD+DCNs under DTX-alpha and individual results superimposed. E- Mean change in the AP repolarizing rate (minimum dV/dt) for all recorded GAD+DCNs under DTX-alpha and individual results superimposed. F- Mean change in minimum ISI-membrane potential (ISI_Vm) for all recorded GAD+DCNs under DTX-alpha and individual results superimposed.

**Figure 2.**
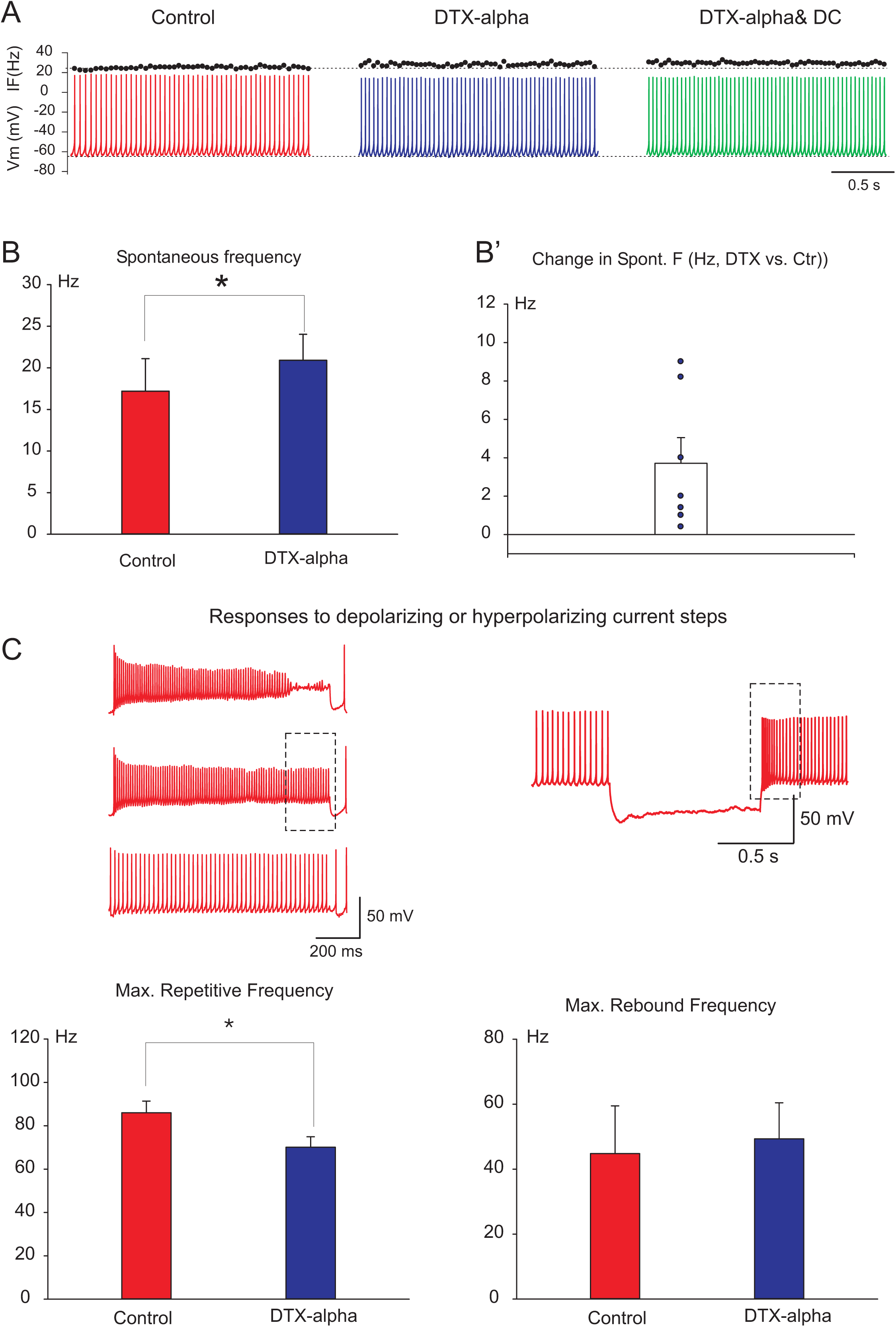
The spontaneous and evoked changes in AP firing rate of GAD+DCNs is sensitive to DTX-alpha. A- From left to right: The traces illustrate typical example of spontaneous action potentials and the dots on top of each spike their corresponding instantaneous frequency using the same scale on the left recorded from one GAD+DCN during application of NT blockers (control), during DTX-alpha (100nM) application (DTX-alpha), application of NT blockers (control), during DTX-alpha (100nM) application (DTX-alpha), and after correcting the ISI Vm by current injection during DTX-alpha application (DTX-alpha& DC) (For comparison, same neuron as in Fig. 1A). B- Mean spontaneous frequency of GAD+DCNs before (control) and during DTX-alpha application (see main text for details). B’- Mean change in spontaneous frequency induced by DTX-alpha application and individual results superimposed (see main text for details). C- Top: left traces illustrate from the typical changes in spiking activity induced by injection of depolarizing current pulses of 1 second duration and increasing current intensity in one GAD+DCN recorded under control conditions used to determine the maximum repetitive frequency (MRF, see methods for details). MRF is determined by measuring the frequency at steady state (rectangle) just before the step inducing depolarizing block (top trace). Bottom left: Mean MRF for GAD+DCNs before (control) and during DTX-alpha application (see main text for details). Top right: The trace illustrates the response induced by injection of hyperpolarizing current pulses of 1 second duration in one GAD+DCN recorded under control conditions. The hyperpolarization typically causes increases in firing rate respect to basal conditions after the hyperpolarization (rebound response). The maximum firing rate over the first second after the step (see methods for details, rectangle, “Maximum Rebound frequency”), was used to characterize changes in the rebound response. Bottom right: Mean Maximum Rebound frequency for GAD+DCNs before (control) and during DTX-alpha application (see main text for details).

Application of DTX-alpha in most cases led to changes in other AP waveform parameters but the results were not consistent amongst neurons and therefore the results were not significant when analyzed as a population. For instance, Kv1 channels have been implicated in some neurons, like pyramidal neocortical cells in the repolarizing phase of action potentials ^31^ and we found that the spike duration, estimated by the duration at half width amplitude (HW), showed increases in some neurons but in other we found decreases (Fig. 1B,top right panel). Changes in the repolarizing rate (estimated by the minimum dV/dt, Fig. 1C, bottom left panel) showed similar variability. Furthermore, the changes in ISI potential (measured without current injection) showed depolarization or hyperpolarization with respect to control levels in different neurons (Fig. 1C, bottom, right panel). Briefly, these results suggest that DTX-alpha sensitive channels control several spike waveform parameters of GAD+DCNs but, that except for the spike threshold, the effects are variable amongst different neurons. The implications will be discussed later.

Next, we investigated the effect of DTX-alpha on the spontaneous action potential firing rate and the changes in firing rate induced by depolarizing and hyperpolarizing current pulse injections (Fig. 2). Application of DTX-alpha led to consistent increases in the spontaneous firing frequency in all neurons tested (n=7). This outcome was independent of whether depolarization or hyperpolarization of the ISI potential was induced by DTX-alpha application. For instance, in the example illustrated in Fig 2A the membrane potential during the ISI hyperpolarized (same example as in Fig.1) and nevertheless the firing rate increased (the instantaneous firing rate is depicted in black on top of the spikes). Constant current injection to bring ISI potential to control levels resulted in even further increase in firing frequency (Fig. 2A, rightmost plot). On average the spontaneous frequency increased by 3.7 ± 1.33 Hz (i.e. 21%, paired t-test, P= 0.0319, n=7. Mean control spontaneous frequency was 17.2 ± 3.9 Hz, Figs. 2B and B’).

DTX-alpha interfered with the ability of GAD+DCNs to fire repetitively at high frequency, tested here using depolarizing current pulses of one second duration and increasing intensity (Fig. 2C, top left plot). The maximum steady state repetitive frequency (MRF, see methods for details) decreased after DTX-alpha application (86 ± 5.4 and 70.1± 4.8 Hz for control and DTX-alpha respectively, n=6, P= 0.031, WSR test, Fig. 2C Bottom left plot). This suggest that the outward Kv1 current is necessary for sustaining high frequency, presumably by enabling sufficient sodium channel de-inactivation between the spikes.

Large DCNs usually respond to hyperpolarizing stimulus with an increase in firing rate above baseline after the stimulus end, so called rebound response e.g.^32, 33, 34^, and GABAergic DCNs also respond to hyperpolarizing inputs with rebound responses ^28^. DTX induced changes in the maximum rebound frequency but we observed increases as well as decreases (max rebound frequency, Figs 2C right plots, see methods) The possible causes of the variability in this or other type of responses will be discussed later.

We investigated as well the effect of other DTXs (I: n=8 and K: n=6) which may have a more specific effect on the Kv1.1 and Kv1.2 channels ^7^ whose mutations are a recognized cause of cerebellar pathology. In general the results using DTX-I or K were similar than with DTX-alpha. After DTX-K application depolarizing block was observed in two neurons, and therefore these were not included in the analysis. Regarding the effect on action potential waveform parameters we found that the more consistent outcome was an hyperpolarization of the AP voltage threshold, measured on averaged action potentials with ISI potential matched to control levels as before, found in all tested neurons except one (the latter showed no difference before and during DTX-I application). Changes in the ISI membrane potential (ISI-Vm) were detected as well but as with DTX-alpha applications these occurred in different directions in different neurons. The action potential duration was slightly increased in all tested neurons, except one where a decrease in HW was found. Regarding the changes in spontaneous firing of action potentials induced by application of DTX-K/I we found increases in the firing rate of most neurons tested (8 out of 10 neurons). Furthermore, application of DTX-K/I led to decreases in the MRF as was the case with DTX-alpha (in 8 out of 10 neurons tested). Regarding the maximum rebound frequency induced after hyperpolarizing current pulses we found diverging effects as with DTX-alpha. Because DTX-K is proposed to be more specific for Kv1.1 channels, the similarity of results using different DTXs supports the idea that the population of channels blocked by the toxins contain at least one kv1.1 subunits ^7^.

Given the similarity of results using these different toxins, we pooled all results together for analysis (Fig. 3, n= 18, The analysed data correspond to GAD+DCNs with long diameters ranging from 7 to 16 μm, mean 12 ± 0.45 μm and the age of the mice from which cells were analysed ranged from P21 to P35, mean P26). On average on the population significant and consistent changes induced by DTXs were found on the action potential membrane threshold, which shifted in the hyperpolarizing direction, (Figs. 3 A and B), the spontaneous firing rate, which increased,(Figs. 3G and H), and the maximum repetitive frequency, which decreased (Figs. 3I and J). DTX application induced changes in the AP duration and its repolarizing rate but the effects consisted in increases or decreases in different neurons. These however, were highly correlated suggesting consistent effects of DTXs in every neuron (Fig. 3F). Application of DTX changed as well the ISI membrane potential in depolarizing or hyperpolarizing direction. Although increases in the spike duration may have caused larger calcium influx and consequently larger activation of calcium-dependent potassium channels controlling the ISI-Vm, overall the changes in ISI potential were not well correlated to those in spike duration (R^2^= 0.034). However, most neurons that hyperpolarized displayed increases in the AP duration (estimated by the duration at half width spike amplitude, HW) during DTX application.

**Figure 3.**
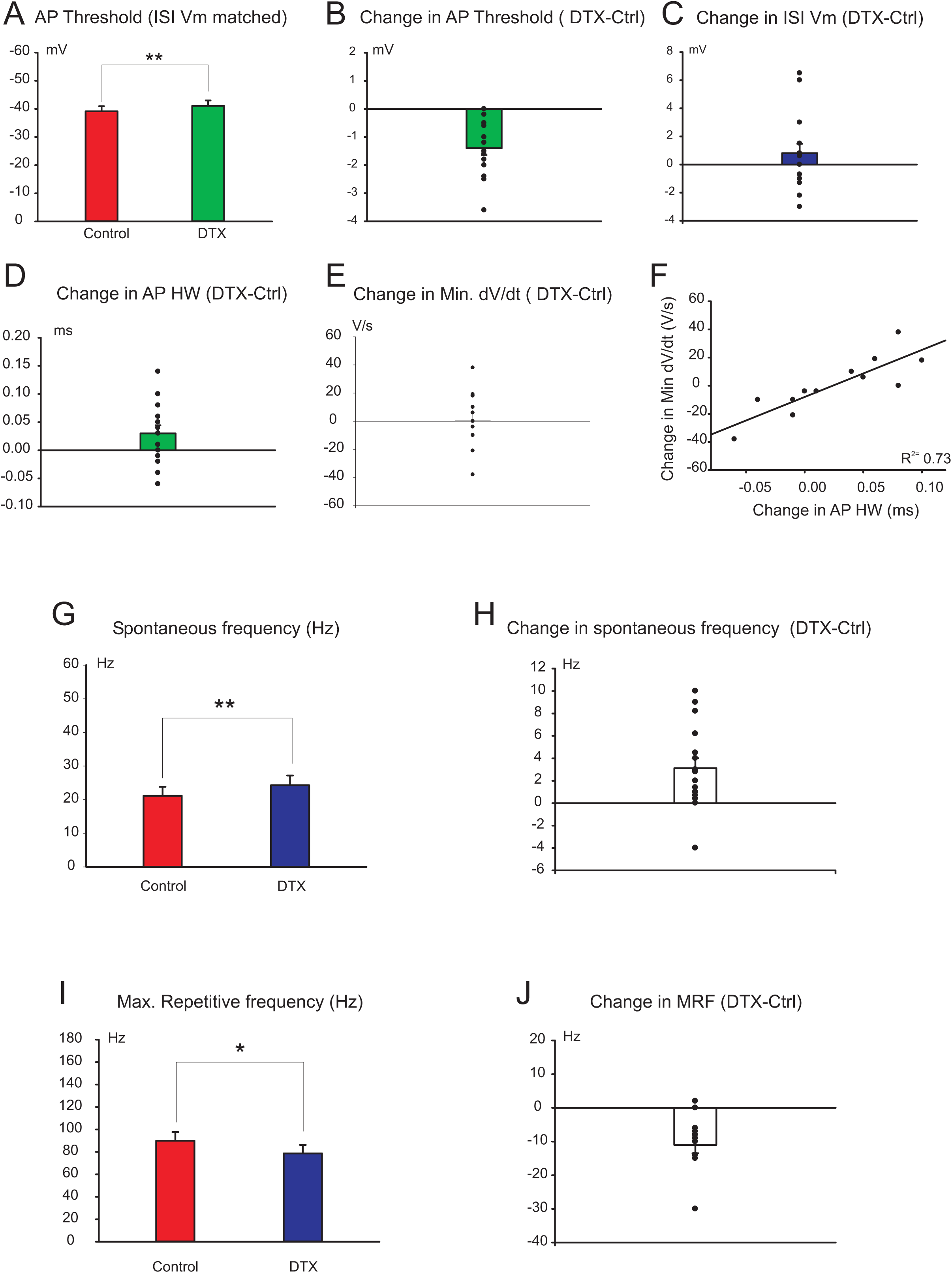
Summary of DTX (pooled DTX-alpha, DTX-I and DTX-K) effect on GAD+DCNs neuronal activity. A-Mean AP threshold under control and DTX conditions: −38.8 ± 1.9 vs. −40.7 ± 2.1 respectively, n=16, paired t-test **P= < 0.001**. B- Mean change in AP threshold and individual results superimposed (same data as in A, - 1.4 ± 0.24 mV). C- Mean change in the ISI-Vm (−0.81 ± 0.67 mV, n= 16, P= 0.247, paired t-test) and individual results superimposed. D- Mean change in the AP-HW (0.026 ± 0.67 ms, n= 14, paired t-test P= 0.109) and individual results superimposed. E- Mean change in the minimum dV/dt (−0.333 ± 5.8 V/s, n= 12, paired t-test P= 0.95) and individual results superimposed. F- Mean spontaneous firing frequency under control and DTX conditions: 21 ± 2.6 vs. 24 ± 2.9 Hz, paired t-test **P= 0.003**, n= 17 G- Mean change in spontaneous firing frequency and individual results superimposed (same data as in F, 3.13 ± 0.89 Hz: 15% of control). H- Mean MRF under control and DTX conditions: 89.9 ± 7.8 vs. 78.7 ± 7.5 Hz, WSR-test **P= < 0.001**, n= 14. I- Mean change in MRF and individual results superimposed (same data as in H, −11 ± 2.5 Hz, 12 % of control).

All these results taken together indicate that GAD+-DCNs activity is sensitive to DTXs strongly suggesting that Kv1 channels regulate these neurons AP waveform and their spontaneous and evoked AP firing activity.

The neurons analyzed so far were GAD+ neurons displaying spontaneous activity. A particular group of GAD+ DCNs co-expressing glycine as neurotransmitter forms a recurrent nucleo-cortical pathway targeting Golgi cells and it has been reported that these neurons are characteristically silent when recorded from slices, in contrast to all other types of DCNs ^24, 26^. During a first series of explorative experiments we tested the effect of the unspecific potassium channel blocker, 4 amynopyridin (4-AP), which at low doses is known to block Kv1 channels (IC_50_ 0.16 to 1.5 mM, ^2^. During these experiments we recorded from two neurons that were silent and therefore could correspond to the subgroup of DCNs forming the nucleo-cortical pathway. In both cases, during application of 4-AP (15μM and 200μM) the membrane potential depolarized and neurons started to fire spontaneously as illustrated in the example of figure 4A. To investigate whether the depolarization was associated to changes in AP waveform as in spontaneously active GAD+DCNs we compared “spontaneous” APs evoked under DTX with those evoked in control conditions by constant current injection or depolarizing current pulses. Indeed, the analysis showed a decrease in the action potential threshold (Fig. 4B, and Fig. 4C), increase in spike amplitude, longer duration mostly due to a slower repolarizing phase and block of the fast AHP. The depolarization and the change in threshold induced by 4-AP application suggested that the “silent” state of these neurons could be under control of the Kv1 channels. However, 4-AP, even at low doses, could block other DCN channels, particularly, Kv3.1 ones ^2, 30^. Therefore, next, we set to explore the effect of DTX on medium sized GAD+-DCNs intrinsically silent. We found only two neurons matching these criteria from a large data set of more than 70 DCNs recorded from the interpositus and lateral nuclei. In these two neurons, application of DTX caused depolarization and induced spontaneous firing. Different than the effect of 4-AP, with DTX (100nM) application no other effect on the AP waveform was detected except for a slight hyperpolarizing shift in the AP threshold (Fig. 4D and E). This suggested that the main effect of Kv1 channels in these neurons is the control of the resting membrane potential and AP threshold while the effect of 4-AP on the repolarizing phase of action potentials is likely due to block of different potassium channels. Overall these results suggested Kv1 channels are key regulators of the state (silent or active) of the GABA/Glycinergic nucleo-cortical pathway neurons.

**Figure 4.**
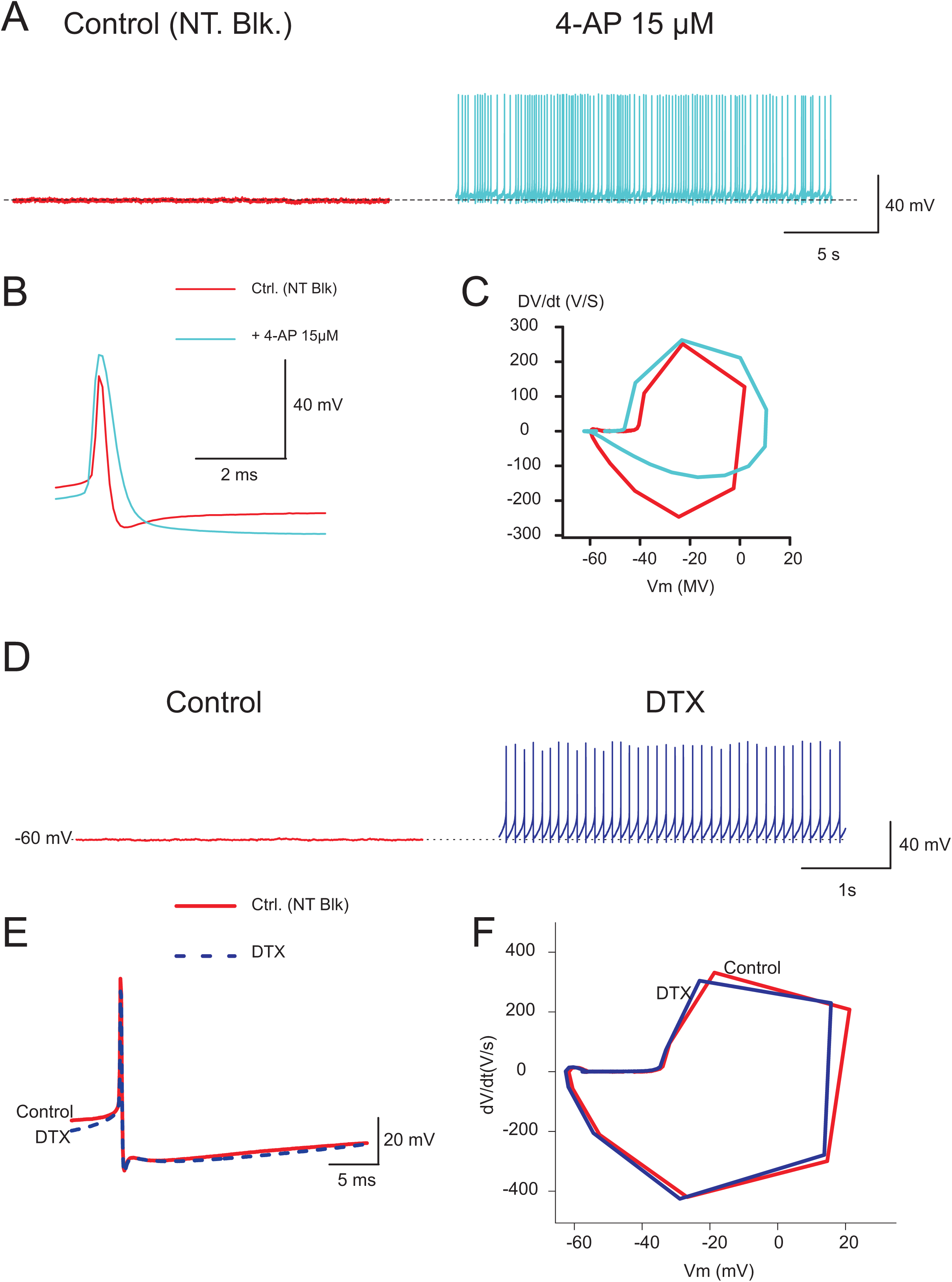
Kv1 channel blockers turn active previously silent DCNs. A- Traces illustrate recordings from one silent DCN before (control) and during application of 4-AP 15μM. Note the depolarization and spiking activity. B- Averaged APs recorded before and during 4-AP application from the same neuron. C- Phase plot corresponding to the traces shown in B. Note the shift in threshold and remarked decrease in minimum dV/dt. D- Traces illustrate recordings from one silent GAD+DCN before (control) and during application of DTX-alpha (100 nM). E- Averaged APs recorded before and during DTX-alpha application from the same neuron as in D. F- Phase plots corresponding to the traces in E.

Finally, to be able to compare the effect of DTX on putatively GABAergic and non-Gabaergic DCNs here we investigated the effect of DTX on putative glutamatergic principal DCNs under the same recording conditions, using animals of the same age and the same type of analysis (Figs 5 and 6). For this set of experiments we used DTX-alpha and DTX-I, and the results were pooled together for analysis.

**Figure 5.**
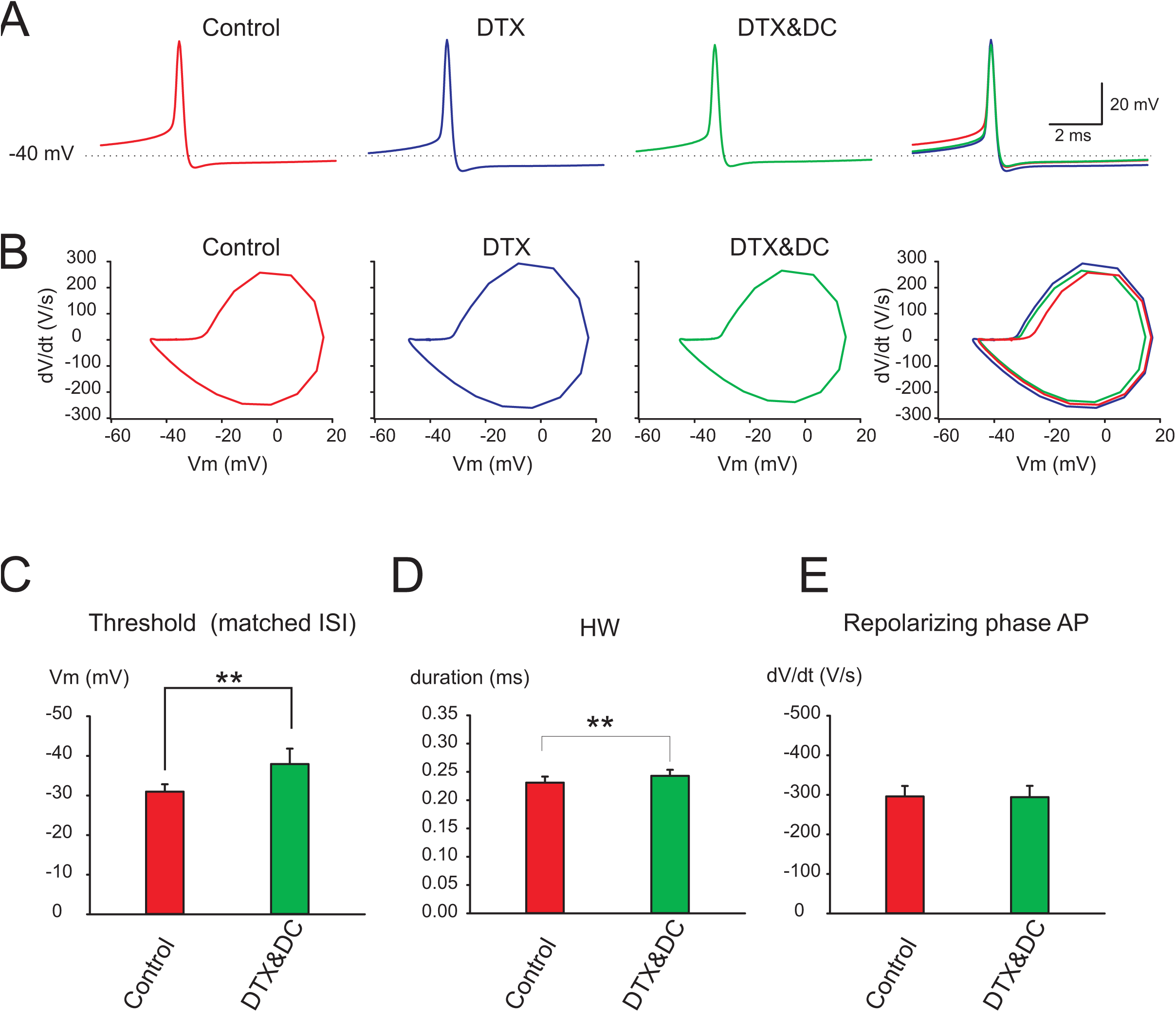
The action potential waveform of non-GAD+DCNs, putative glutamatergic principal DCNs is sensitive to DTX. A- From left to right: Typical example of averaged spontaneous action potentials recorded from a large Non-GAD+DCN during application of NT blockers (control), during DTX-alpha (100 nM) application (DTX-alpha), and after matching the ISI Vm to control levels by current injection during DTX-alpha application (DTX-alpha& DC), and all traces superimposed. B- The corresponding phase plots of the traces illustrated in A. C- Mean AP threshold before and during DTX application for putative glutamatergic principal DCNs (see text for details). D- Mean AP HW before and during DTX application for putative glutamatergic principal DCNs (see text for details). E- Mean AP repolarizing rate (minimum dV/dt) before and during DTX application for putative glutamatergic principal DCNs (see text for details).

**Figure 6.**
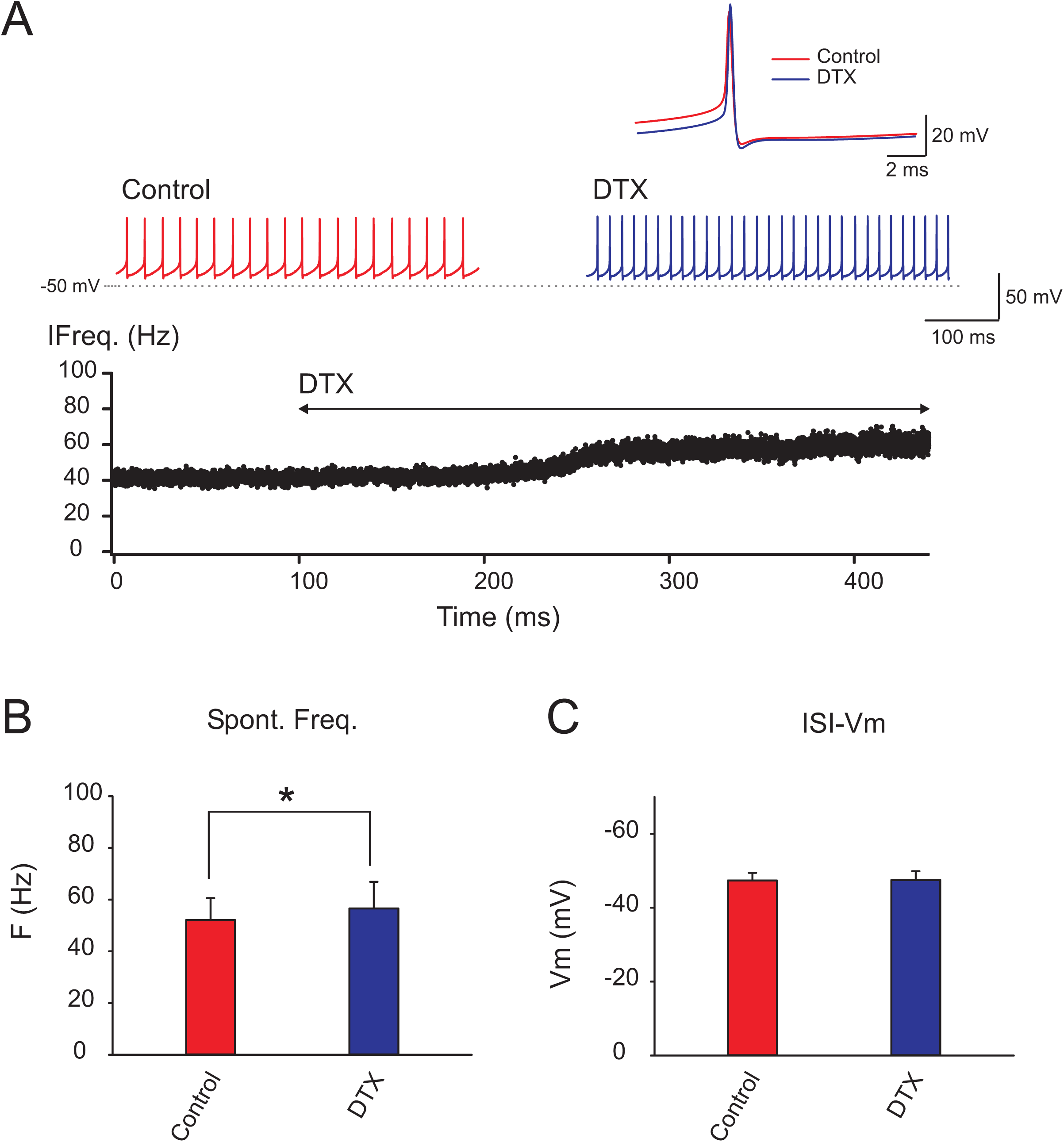
Kv1 channels control the spontaneous firing rate of non-GAD+DCNs, putative glutamatergic principal DCNs. A- The red and blue traces illustrate segments of the whole cell current clamp recordings of a spontaneously active putative glutamatergic principal DCNs before and during application of DTX-I (100nM) respectively. Note the depolarization and decrease in ISI between APs during DTX application. The bottom plot illustrates the changes in instantaneous frequency induced by application of DTX to the bath. The inset above shows the averages of APs obtained before and during DTX application (red and blue, respectively). B- Mean spontaneous frequency before and during DTX application for putative glutamatergic principal DCNs (see text for details). C- Mean ISI membrane potential before and during DTX application for putative glutamatergic principal DCNs (see text for details).

Similar to the case with GAD+DCNs, and consistent with previous results ^17^, DTX modified the activity of hthese neurons. A change in the AP membrane threshold was the most consistent effect of DTX on the AP (Fig 5A, B and C). On average the threshold hyperpolarized 3.3 ± 0.83 mV (n=10, paired t-test, P= 0.003). The AP duration estimated by the HW was slightly increased under DTX application (5 %, 0.23 ± 0.011 and 0.24 ± 0.034, paired t-test P= 0.009, n= 10). However, no significantly changes in the repolarizing rate were detected (minimum dV/dt, paired t-test P= 0.873, n=10).

Application of DTX led in most neurons to increases in spontaneous firing rate (Fig. 6, on average 21%: 48.5 ± 7.9 and 58.9 ± 10.8 Hz for control and DTX respectively, paired t-test P= 0.028, n=10). In addition, similar to GAD+DCNs, DTX application decreased the ability to fire repetitively at high frequencies (estimated by the MRF, 441 ± 33.4 and 412 ± 28 Hz for control and DTX respectively, WSR test P= 0.023 n=9). The ISI membrane potential depolarized or hyperpolarized in different neurons similar to the effect of DTX on GAD+DCNs. Summarizing, the main effect of DTX on putative principal DCNs was a decrease in AP threshold, increase in spontaneous firing rate and decreased ability to fire at high frequencies.

## DISCUSSION

Here we used a pharmacological approach to investigate the role of low threshold potassium currents mediated by Kv1 channels in regulating GAD+DCNs, putative GABAergic DCNs. The results indicate that the activity of this group of neurons is sensitive to DTX-alpha, DTX-I and DTX-K. In particular we observed consistent and significant hyperpolarization of the action potential (AP) membrane threshold, increase in the spontaneous firing rate and decreases in the maximum frequency at which neurons could sustain repetitive firing. Furthermore, we confirmed that putative glutamatergic principal DCNs are sensitive to DTXs: these responded with similar hyperpolarization of action potential threshold, increase in spontaneous frequency and decrease in the ability to fire repetitively at high frequency. Finally, in a group of medium sized DCNs silent in control conditions, application of Kv1 channel blockers induced depolarization and tonic firing of action potentials. Overall, these results strongly suggest that Kv1 currents stabilize the AP threshold and spontaneous pacemaker activity of GABAergic-DCNs, while facilitating the high frequency responses to excitatory inputs.

Kv1 channel have low voltage activation ranges and therefore are suited to control the AP threshold and neurons excitability as demonstrated in other neurons^2, 3, 31^. The spike voltage threshold is assumed to result from the interplay between the inward sodium current and the outward currents active around the point in space and time where the sodium current becomes regenerative ^1^. DTX shifted in the hyperpolarizing directions the voltage threshold of spontaneous APs although the spontaneous firing rate increased. The increases in firing rate were associated to decreases in spike amplitude and AP depolarizing rate (estimated by the dV/dt) and this effect was larger when depolarizing constant current was injected to match the control membrane potential (e.g. Fig. 1A DTX&DC). This suggested that increases in frequency shortens the available time for sodium channel de-inactivation between the spikes and therefore the proportion of sodium channels available for the next spike. Although enhanced inactivation of sodium channels could have shifted the voltage threshold in the depolarizing direction ^35^, the spike voltage threshold hyperpolarized under DTX, suggesting a robust role for Kv1 channels in controlling this parameter. In other neurons it has been demonstrated that local clustering of Kv1 channels around the site of AP initiation is critical for their control of the voltage threshold. Further studies could elucidate if this is the case for DCNs.

DCNs are spontaneously active and the membrane potential between spikes is relatively depolarized and therefore within the activation range of Kv1 channels ^2, 4, 7^. Our results indicate that the spontaneous activity and firing rate is sensitive to Kv1 channel blockers suggesting that Kv1 channels continuously control the output of GABAergic (as well as non-GABAergic) DCNs. In part this effect can be attributed to the shift in AP threshold. Indeed, repolarization to control ISI potential with current injection in the case of cells that depolarized was still associated to increased frequency, supporting this view. In addition, at least in some neurons, direct depolarization of the ISI membrane potential could have contributed to the increased frequency.

Our recordings likely include different types of GABAergic-DCNs, including the nucleo-olivary cells, the GABAergic interneurons, and perhaps the nucleo-cortical GAD+DCNs. By controlling the spontaneous tonic firing rate of these different GABAergic DCNs, kv1 channels could be in control of 1) the occurrence of Purkinje cells complex spikes by controlling the nucleo-olivary pathway activity. 2) the cerebellar output: DTX application increased the activity of the principal DCNs, but at the same time increased the activity of GABAergic DCNs, suggesting opposing effects of the same channels on this output. An interesting question is how these two opposing effects interact under different conditions. 3)Finally our results indicate that the group of silent DCNs, which are proposed to mediate a nucleo-cortical inhibitory pathway controlling cerebellar input stage ^24^, have their silent state under kv1 channels control. Changes in the tonic level of activity of these neurons could have a large impact on the gating effect of Golgi cells and therefore on the activity of granule cells inputs to Purkinje cells. Furthermore depolarization of these neurons could compromise their ability to fire bursts of action potentials and therefore their usual function^24^. We found only a small number of silent DCNs. However, one possible explanations is that we fail to detect larger numbers because GAD-GFP fluorescence is lower than average at these cells, or that these cells at close to physiological temperatures are not silent. In any case, further studies should address this issue.

Other parameters of the AP waveform were affected by DTX application, but the direction of changes and magnitudes were variable. Part of these effects could be directly driven by the block of Kv1 channels, while other could be indirectly caused by them. For instance, the hyperpolarization of the ISI Vm is unlikely to be caused directly by the block of Kv1 channels, which mediate low threshold outward currents ^4^. Instead, elongation of AP by interference of Kv1 channels with the AP repolarization could have caused increase in calcium dependent potassium currents responsible for the hyperpolarization between spikes. Kv1 channels have been implicated in some neurons in the repolarizing phase of APs. However, the increases in DCN spike duration caused by DTX were only modest in the cases that occurred (maximum was 16%), in contrast to those attributed to Kv3 channels 30. This suggests that differences in the proportion of different currents supporting AP repolarization in each DCN could explain the variable effects of DTX. Similar arguments can be attributed to the effect of DTX on the membrane potential of spontaneously active DCNs: differential contribution of membrane channels to the ISI potential could explain different effects. Instead, DTX induced a clear depolarization on the group of silent DCNs, suggesting a bigger role for these channels in the control of the resting membrane potential for this group of neurons.

The differences in responses to DTX could be explained by the presence of different GAD+DCN classes. At least two different subgroups, the nucleo-olivary DCNs and the GABA/glycine interneurons that innervate the DCNs principal cells compose spontaneously active GABAergic DCNs. Although the reported electrophysiological properties of the two groups of neurons cannot separate them apart ^28, 36^ it could be that non-evident differences in the currents underlying their spontaneous firing could explain heterogeneity in DTX responses. Alternatively but not excludingly, it could be that each neuron expresses a range of different proportion of currents independently of whether each is part of a particular class, as has been proposed for other systems ^37^.

Taken together these results indicate that spontaneous activity and responsiveness of GABAergic DCNs is controlled by Kv1 channels. From a functional point of view, and since there is evidence that Kv1 channel expression can be modulated by synaptic activity^38^, it will be interesting to investigate the possibility of regulation of Kv1 channels at GABAergic DCNs and whether this modulates spontaneous and responsive properties of these neurons and thereby diverse aspects of cerebellar function. From a pathophysiological point of view, these results point to a new cellular cause for cerebellar dysfunction in forms of episodic ataxia involving Kv1 channel mutations, and potential new avenues for their treatment.

## Acknowledgements

I thank Cornelius Schwarz for support for this research, alongside that from the Cognitive Neurology Department; Ute Grosshennig and Ursula Pascht for technical assistance; all members of the Cognitive Neurology Department for comments and helpful discussions.

## METHODS

The animal protocol was reviewed and approved by an independent local committee and the Regional Council of Tübingen and conducted according to the standards of German law and the Society for Neurosciences (SFN).

### Cerebellar slices preparation

For preparing cerebellar slices mice from the GAD67-GFP knock-in mice line (with one GFP protein gene substitutes one GAD-67 allele) ^39^ their, wild type littermates or wild type animal with the same background (either sex, >P21) deeply anesthetized with ketamine (150 mg/kg) or flurazepan were prepared as previously ^33^using a vibrotome (Leica, Bensheim, Germany) and ACSF (containing in mM: 125,5 NaCl, 2,5 KCl, 1.3 NaH_2_PO_4_, 1 MgCl_2_, 26 NaHCO_3_, 20 Glucose, 1.5 CaCl_2_), bubbled with 95% O_2_ and 5% CO_2_ maintained at 26°C. Slices were immediately transfer to a storing chamber with ACSF at 36°C and afterwards let to cool down to room temperature in the same solution until recorded. The same solution at close to physiological temperatures (33-35 °C) was used for recordings using a submerged type chamber.

### Whole cell Patch clamp recordings

Whole-cell current clamp recordings were made from DCNs in the lateral or interpositus nuclei using an Axoclamp2B-amplifier (Molecular Devices). DCNs were visualized using infrared illumination and the presence of fluorescence was tested using LED illumination (470nm, CoolLED).The intracellular solution contained (in mM): 134 Kgluconate, 6 KCl, 10 KHEPES; 0.1 EGTA, 0.3 NaGTP, 2 KATP, 10 Phosphocreatine, 2 MgCl_2_. Recordings were digitized (12.5 kHz), and stored using programmable software (Spike 2, CED) for further analysis.

### Drugs

The following drugs: gabazine (3μM) an antagonist of GABAA receptors, strychnine (1μM) an antagonist of glycine receptors, and kynurenic acid (3-5mM) a broad-spectrum antagonist of ionotropic glutamate receptors were systematically applied to the ACSF to be able to isolate pharmacologically the effect of the Kv1 blockers on the intrinsic properties of DCNs (Control conditions). (All these antagonists were purchased from Sigma). In a series of explorative experiments, 4-aminopyridine (15 or 200μM), a broad spectrum potassium channel blocker was added to the bath contanining neurotransmitter blockers. Dendrotoxins DTX-alpha, DTX-I and DTX-K (100nM) were diluted as recommend by the providers (Alomone, Latoxan) to prepare aliquots that were frozen and diluted to the final concentration in ACSF containing neurotransmitter blockers before use. We noticed that the potency of the toxins decreased with time after aliquot preparation. Therefore, when this information was available, we excluded experiments using aliquots older than 4 weeks.

### Experimental protocols and analysis

protocols and analysis were done as previously ^30, 34^. Briefly: the recordings were performed at the spontaneous membrane potential unless otherwise noted. Only neurons with stable recordings were included for analysis. Rarely, during recordings some neurons “spontaneously” depolarized and remained in depolarized state (depolarizing block). These neurons were not included in the analysis although sometimes they recover. Because one effect of the drug application could be to facilitate this process, we report the occurrences after toxin application, but we did not include data from these neurons in the analysis. For the spike waveform analysis of action potentials (APs) we used averages of successive spikes aligned by their peak. The threshold was detected using the time derivative of the membrane potential in the phase plots, as the point where the membrane derivative sharply rose or the values were higher than 5-10 V/s. The AP amplitude was taken as the difference in potential between the threshold and the AP peak. The AP half width was the duration of the spike at half amplitude. To estimate changes in the rate of the depolarizing and repolarizing AP phases the maximum and minimum values of the AP derivative were used respectively.

The basal AP firing rate was calculated for periods of at least 10 seconds of stable basal activity.To emulate synaptic evoked activity, hyperpolarizing or depolarizin current pulses (1s duration) were injected. The maximum rebound frequency, was defined as the maximum instantaneous frequency attained over the first second following the end of hyperpolarizing pulse injections of increasing intensity. To evaluate changes in the ability to fire repetitively at high frequency, the maximum repetitive firing frequency (MRF) was defined as the maximum steady state frequency during the last 200 ms of the responses to depolarizing current pulses of increasing intensity. Analysis was performed using programmable software: (Spike 2, CED), Igor (Wavemetrics Inc.) and further statistical analysis was carried out using Sigma Plot (SPSS Inc.). Data are presented as means ± SEM, sample size and type of statistical test used to assess significance are indicated in the main text and/or figure legends.

